# Microtubules and Gαo-signaling independently regulate the preferential secretion of newly synthesized insulin granules in pancreatic islet β cells

**DOI:** 10.1101/2020.10.26.354936

**Authors:** Ruiying Hu, Xiaodong Zhu, Mingyang Yuan, Kung-Hsien Ho, Irina Kaverina, Guoqiang Gu

## Abstract

For sustainable function, each pancreatic islet β cell maintains thousands of insulin granules (IGs) at all times. Glucose stimulation induces the secretion of a small portion of these IGs and simultaneously triggers IG biosynthesis to sustain this stock. The failure of these processes, often induced by sustained high-insulin output, results in type 2 diabetes. Intriguingly, newly synthesized IGs are more likely secreted during glucose-stimulated insulin secretion. The older IGs tend to lose releasability and be degraded, which represents a futile metabolic load that can sensitize β cells to workload-induced dysfunction and even death. Here, we examine the factor(s) that allows the preferential secretion of younger IGs. We show that β cells without either microtubules (MTs) or Gαo signaling secrete a bigger portion of older IGs, which is associated with increased IG docking on plasma membrane. Yet Gαo inactivation does not alter the β-cell MT network. These findings suggest that Gαo and MT regulate the preferential release of newer IGs via parallel pathways and provide two potential models to further explore the underlying mechanisms and physiological significance of this regulation in functional β cells.

## Introduction

In response to postprandial blood-glucose increase, pancreatic islet β cells secrete insulin to promote glucose usage and storage in peripheral tissues (the liver, fat, and skeletal muscle), ensuring blood-glucose homeostasis. The collective β-cell dysfunction, loss-of identity, or death (a.k.a. β-cell failure) results in inadequate insulin secretion (1–3). This leads to type 2 diabetes (T2D), featured by sustained high blood-glucose levels and deregulated lipid metabolism that damage multiple tissues (4). In contrast, excessive insulin secretion, caused by cancerous β-cell proliferation (5) or deregulated secretion (6, 7), results in hyperinsulinemic hypoglycemia that leads to comatose or even death.

To control insulin secretion, β cells precisely regulate insulin biosynthesis, storage, transport, and secretion (8). Each β cell contains around 10,000 IGs (9). According to the physical location of and response to stimulus, these IGs have been traditionally classified into two pools: the readily releasable pool (RRP) and reserve pool (RP) (10). The former refers to a small group of IGs (<5%) that are docked onto the plasma membrane (PM) (11). These IGs were immediately released upon stimulation and largely contribute to the first phase of insulin secretion (11). In dysfunctional islets from T2D patients, β cells lack this IG pool, so that they cannot quickly secrete insulin in response to glucose stimulation within the first few minutes, i. e., the first phase (12). The RRP is absent in newly differentiated immature β cells as well, likely depleted by high levels of basal secretion (13).

The RP contains the majority of IGs in β cells, which are usually located away from the PM. These vesicles need transport to the PM for docking and priming to be released. To this end, high glucose, besides triggering glucose-stimulated insulin secretion (GSIS) and new IG biosynthesis, induces the transport and conversion of some IGs from RP to replenish the RRP (10, 14–17). These inter-connected responses allow β cells to maintain sustained or pulsatile GSIS under continuous or pulses of glucose stimulation, a property that is necessary for long-term β-cell function.

Intriguingly, not all IGs in the RP are alike and are able to be mobilized to the RRP. Several studies, using pulsed radio-labeling or fluorescent-protein tagging of insulin, have shown that newly synthesized IGs are more likely released upon stimulation (18–23). Aged IGs will become non-functional and degrade via proteolysis (24, 25). This degradation ensures long-term β-cell function by removing the non-responsive IGs. It also presents additional metabolic load due to the futile biosynthesis of these IGs, which contributes to the high β-cell stress and reduced cell proliferation/function (26–29). Thus, investigating how β cells preferentially secrete newly synthesized IGs can lead to ways to enhance insulin output while avoiding insulin biosynthesis-induced dysfunction.

A feature that potentially contributes to the preferential release of new IGs is their transportability via the microtubule (MT) network. In an elegant study of temporally-marked IGs, Hoboth and colleagues showed that IGs can display three types of glucose-modulated and MT-dependent mobility: highly dynamic, restricted, and nearly immobile states. High glucose can expedite this motion, with the young IGs being more responsive to glucose, and older ones less sensitive and more likely found in the lysosome (30). These findings support a model that old IGs tend to lose MT-dependent transportability, preventing their movement to the PM and reducing their chance of docking/release.

MTs are cytoskeletal biopolymers that act as tracks for vesicular transport using the kinesin and dynein motor proteins (31, 32). In many cell types, MTs originate from the centrosome to form a radial array, with their plus ends oriented toward the cell periphery. Henceforth, kinesin- or dynein-mediated transport mediates the bulk flow of cargo toward the cell periphery or interior, respectively (33). In contrast, most of the β-cell MTs originate from organizing centers in the Golgi and form a non-directional meshwork (34, 35). This network is essential for quick/long-range IG movement but is ill-suited for bulk directional cargo transport (34–39). Disrupting these MTs acutely enhances GSIS (34, 35, 40), while stabilizing the MTs represses secretion (34, 35, 41). A model that is supported by these findings is that β-cell MTs allow active IG-movement between cell interior and cell periphery; however, MTs also compete with the PM for IG binding to acutely reduce the RRP. In the absence of MTs, the overall long-range IG transport is reduced (35, 38, 39). Yet the non-MT-dependent IG movement, within a brief period of time, is sufficient for a portion of IGs to move to the PM for docking and regulated release (35). Thus, regulating the dynamics and density of MTs will likely influence the releasability of young versus old IGs, because the new IGs will lose their advantage of being moved to cell periphery when MT-aided transport is reduced.

In addition to transport, IG docking on the PM is another limiting step for insulin secretion (15, 17, 42, 43). In this case, vesicular and PM proteins form a SNARE complex via the association between Synaptobrevins, Syntaxins, SNAP23/25, Munc18, Rim, and others (44). This complex brings the vesicles close to the PM. The presence of Ca^2+^, via a family of Ca^2+^ sensors such as Synaptotagmins (Syts) (13, 45, 46) and/or Doc2B (47–49), further modulate the conformation of the SNARE-complex to enable vesicular/PM fusion (50). Thus, mutations in several of these SNARE components were found to deregulate IG docking and secretion, and underscore essential roles of docking in GSIS (17, 43, 51–54).

An intriguing signaling molecule that can potentially regulate both IG docking and MT dynamics is inhibitory G protein Gαo. Like all other Gα subunits, Gαo signaling toggles between on- and off-state by dissociating/associating with Gβγ dimers in response to the activation of G-protein coupled receptors (55). Unlike other inhibitory Gα (Gαi1, Gαi2, Gαi3, and Gαz), Gαo activation does not inhibit adenylyl cyclases in several cell types including islet β cells (15, 56), but repressing GSIS by reducing IG docking (15). Intriguingly, Gαo at high levels, together with other inhibitory Gα subunits, was shown to promote MT disassembly by Gα-MT association (57, 58). Here, we explore the hypothesis that Gαo may regulate IG transport/docking through the MT network, which consequently control the probability of IG secretion in young and aged IG pools.

## Results and discussion

### The β-cell MTs are dispensable for overall IG-distribution to cell periphery

We have previously shown that the MT network in β cells, although essential for sustained secretory function (40), acutely represses GSIS (35, 40, 59). Based on Total Internal Reflection Fluorescence Microscopy (TIRFM)-observation of MT-dependent IG movement near cell membrane (<200 nm away from the PM), we postulated that a role of the MT network in β cells is to pull IGs away from the PM besides acting as tracks for long-distance IG movement (34, 35). Here, we corroborated this hypothesis by first examining the overall distribution of IGs in β cells with or without MTs.

Isolated islets were stained for insulin immunofluorescence (IF) and confocal microscopy after incubation in 10 μg/ml Nocodazole (NOC) for 12 hours, a condition that effectively destabilize MTs (Figure 1A, B). We then examined the insulin levels at different areas of β cells, starting from one spot of β-cell membrane to another on the opposite side of cell membrane (see white lines in Figure 1C). At both non-stimulating 2.8 mM glucose (G2.8) and stimulating G20, there were significant reductions in total IG levels when cells were treated with NOC (Figure 1D), with p = 0.0015 and p<0.0001 (ANOVA), respectively. We observed particularly high IG reduction in the center of cells in islets treated with G20 and NOC, when the overall IG was quantified (Figure 1D) or when the relative distribution of IGs (i.e., the % of IGs distributed at each spot of β cells) were examined (Figure 1E). These findings are consistent with our conclusion that MTs are dispensable for the movement of IGs to cell periphery, likely achieved by slow-but-detectable random IG movement or MT-independent transport (35).

**Figure 1.**
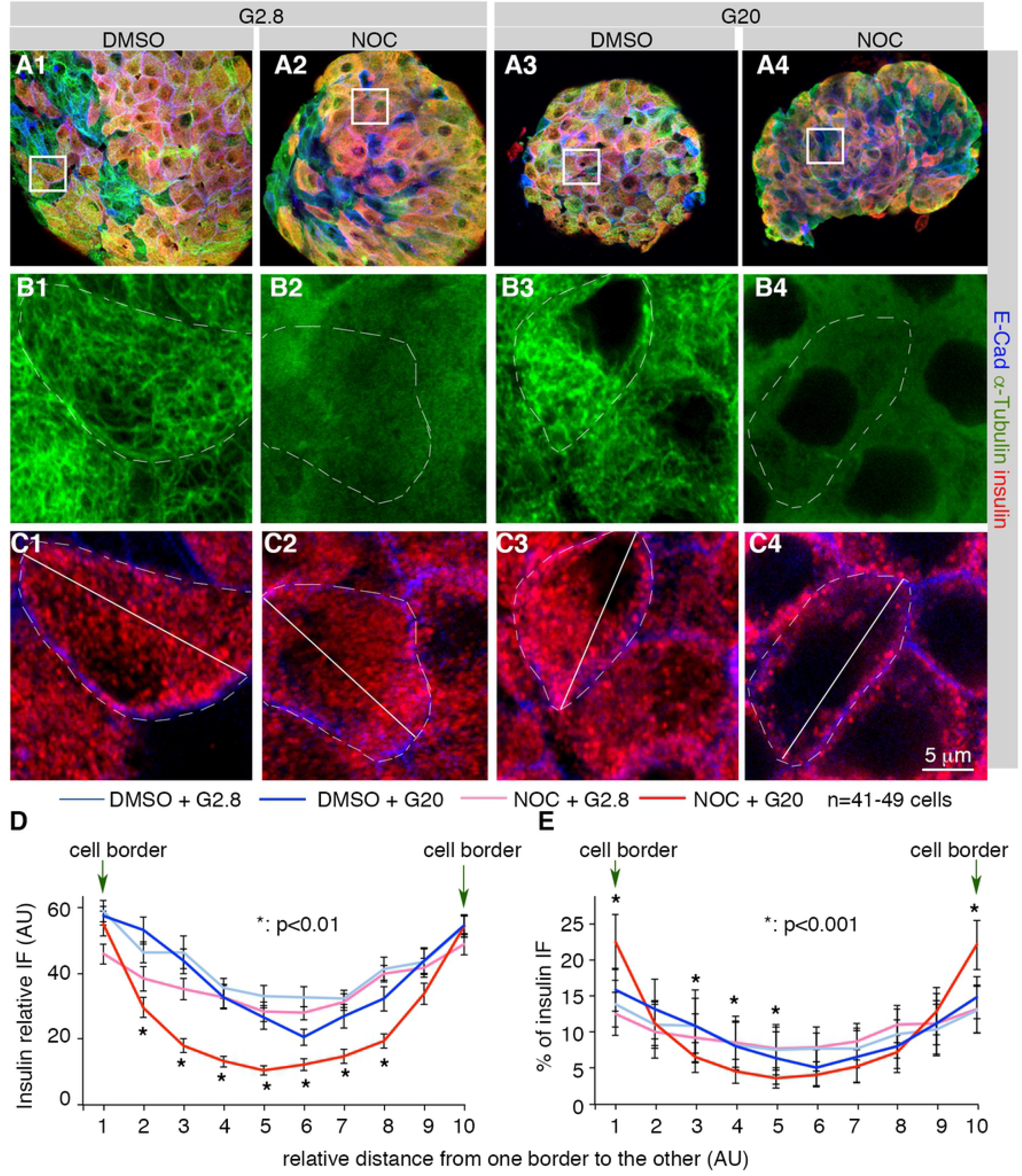
MTs are dispensable for the allocation of IGs near the β-cell periphery. (A-C) IF-assays of IG distribution in intact islets treated with [2.8 mM glucose (G2.8) + DMSO] (column 1), (G2.8 + NOC) (column 2), (G20 + DMSO) (column 3), and (G20 + NOC) (column 4). Images shown in (A) are maxi-projections of Z-stacked optical sections, showing co-staining of insulin (red), E-cadherin (E-Cad, blue), and α-tubulin (green). (B, C) high-magnification images showing α-tubulin (B) or (insulin + Ecad, C) IF signals in boxed areas of (A). Broken white circles highlight quantified β cells. Panel (B) images verify the absence of detectable MT filaments with NOC-treatment. Panel (C) shows the way of quantifying insulin distribution - with a line-scan along the white line drawn across each β cell. (D) IF intensity profile of insulin along the long axis of β-cells from confocal images. (E) Another way to show the insulin-distribution patterns with/without MT disruption, with the % of insulin detected at different locations in β cells along the long axis. P values marked in both (D) and (E) were calculated with two-tailed type II t-tests from the data of (DMSO + G20) and (NOC + G20) groups.

### The β-cell MT inhibits IG docking on PM

We next examined the number of docked IGs via TEM in the absence of MTs, focusing on vesicles <10 nm away from the PM, a resolution that cannot be achieved by TIRFM or confocal microscopy (Figure 2A, B). MT-depolymerization significantly increased the number of IGs docked onto the β-cell PM at basal glucose (Figure 2C). In contrast, there is a significant decrease in the density of IGs in β cells without MTs (Figure 2D-F), consistent with the requirement of MTs for new IG biosynthesis (40). Similarly, the increased IG docking was observed in the absence of MTs under high glucose conditions (Figure 2G-I), despite significant degranulation with or without MTs under high glucose (Figure 2J. Compare with Figure 2F).

**Figure 2.**
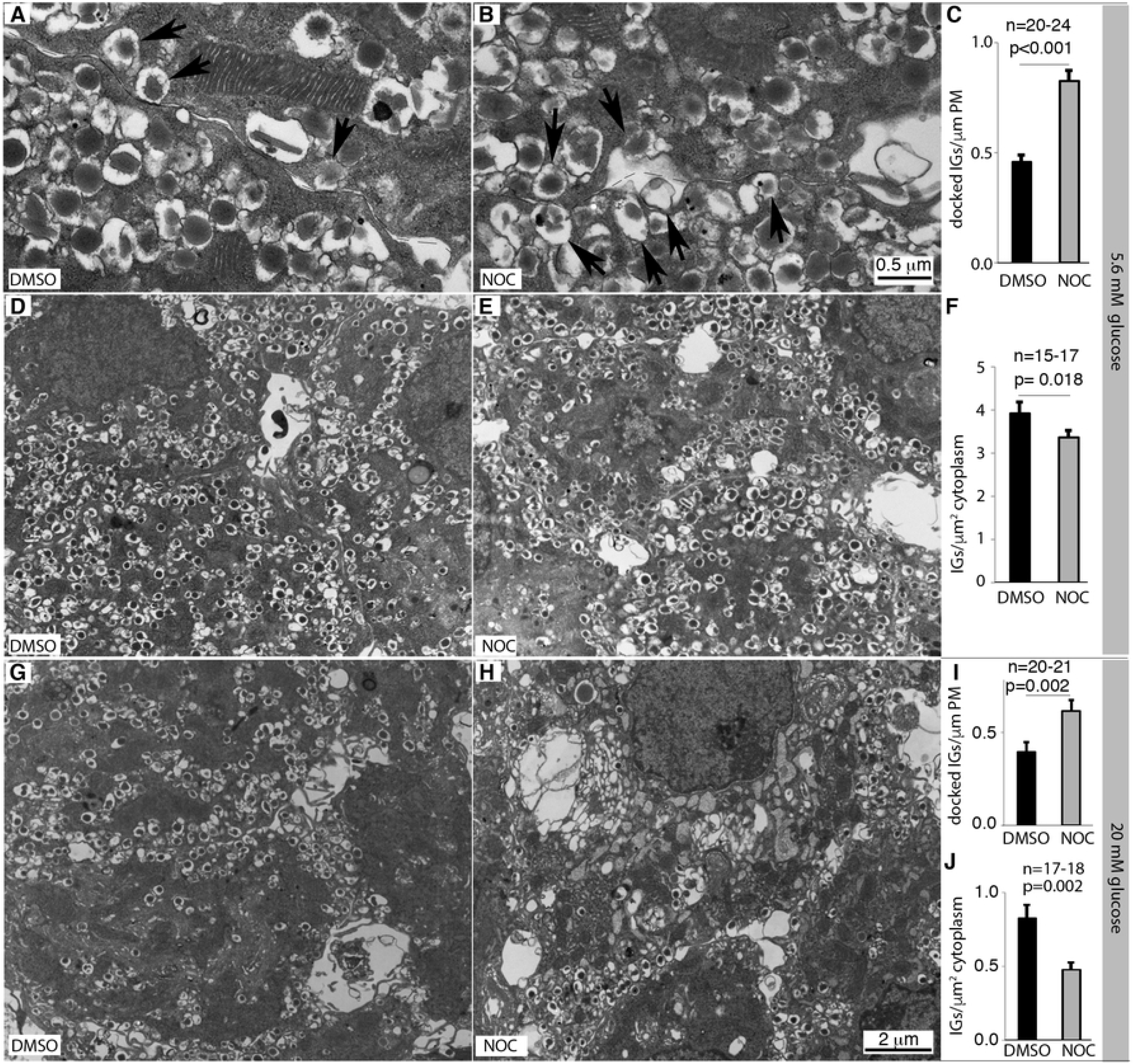
TEM imaging showing the inhibitory effects of MTs on IG-docking in β cells. Islets were isolated from wild-type (WT) adult ICR mice and were treated with DMSO or 10 μg/ml NOC for ~12 hour at 5.6 or 20 mM glucose. TEM was then used to examine the locations and density of IGs. (A-C) TEM images and quantification of docked IGs from DMSO- and NOC-treated islets in the presence of 5.6 mM glucose. (D-F). Images and quantification of IG density in microscopic fields used in (A-C). (G-J) Images and IG quantification as in (A-F), except 20 mM glucose was used. Scales in (D, E, G, H) are the same, labeled in panel (H). In (C), (F), (I), and (J), mean + SEM were presented. In all panels, “n” indicates the number of microscopic fields (with 3-4 different β cells included in each field) counted. P, results from two-tailed type II t-test.

These results are consistent with a model that MTs compete with the PM for IG binding. Specifically, IGs likely associate with MTs via vesicle-bound motor proteins. This allows the MT-dependent IG-transport from the site of biogenesis, the trans-Golgi network that usually localizes in interior of cells, to close-to the PM (34, 36–39, 60, 61). However, the β-cell MTs have no obvious directionality. Thus, the MTs can also pull IGs away from the PM to prevent IG docking and to reduce the RRP, supported by both experimental results (17, 35) and mathematical modeling (34). When MTs are destabilized near the PM, e.g., in the presence of high glucose, the IGs can lose MT contact and be available for docking/secretion. Consequently, IGs with preferential binding with MTs, especially those newly synthesized (30), are more likely transported to the cell periphery for docking and release (59). The older vesicles, less likely transported due to their attenuated MT-association, will eventually be degraded.

The above model predicts that when the MT network is disrupted, the fast IG movement will be abolished (30, 36). Yet vesicle movement [slowed but still detectable in the absence of MTs (35)] via free-diffusion or actin-assisted transport is probably sufficient for a portion of IGs to move to underneath the PM for docking and regulated secretion (58, 62). In this setting, older IGs can be mobilized to allow enhanced secretion without new IG biosynthesis. In addition, the newly synthesized IGs with superior MT-dependent transportability will lose the advantage of being moved to underneath the PM (30). In other words, the absence of MTs could normalize the probability of secretion for both new and older IGs, which we experimentally evaluated next.

### Disrupting the MTs allows increased secretion of older IGs from β cells

To test if MT-disruption induces increased secretion of old IGs, GSIS was carried out in the absence of protein synthesis and MTs. Islets were incubated in the presence of 10 μM cycloheximide (CHX) for three-hours, a condition that can effectively (>95%) inhibit protein biosynthesis in islets (63). Treated islets were then incubated with 10 μg/ml NOC for one more hour to disrupt islet-cell MTs (Figure 3A-D), followed by GSIS assays. Because newly synthesized insulin become secretable within two-hours (64), we expect this treatment to reduce newly produced secretable IGs. The lack of protein biosynthesis results in a significant reduction in GSIS in β cells with functional MT (i.e., without NOC-treatment) (p= 0.02, Figure 3E) as expected (63). Yet this lack of protein synthesis did not eliminate the MT-destabilization-enhanced GSIS (Figure 3E).

**Figure 3.**
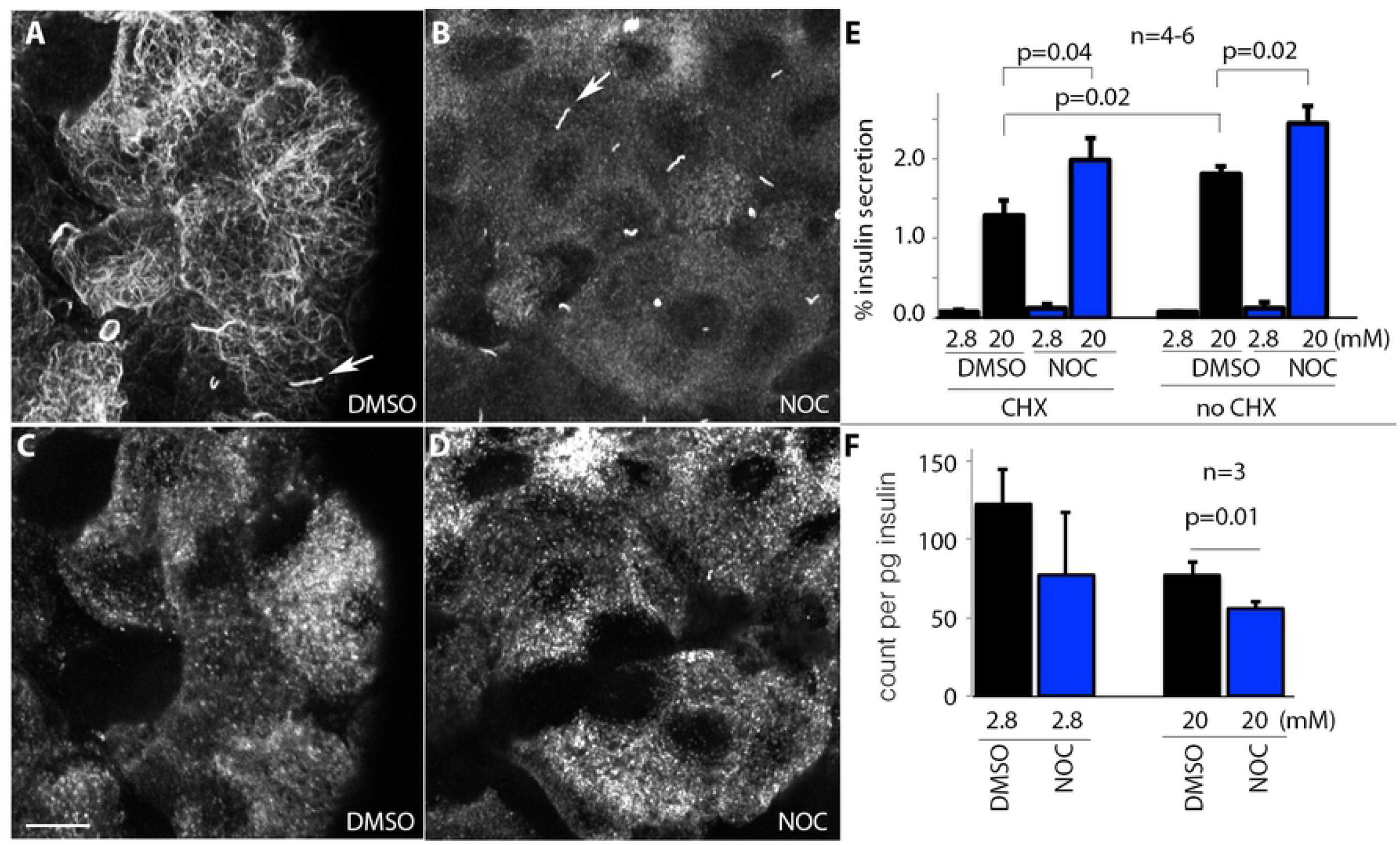
The absence of MTs promotes secretion of aged IGs. (A-D) The effective MT-disruption in islet β cells with NOC. Glu-tubulin was stained (A, B) to verify the effective disruption of MTs by NOC, with β cells identified by insulin staining (C, D, corresponding to the fields in A, B, respectively). Arrows in (A, B), primary cilia. Scale bar = 10 μm. (E) GSIS from islets treated with combination of 10 μg/ml NOC and 10 μM CHX. (F) The relative amount of radioactive insulin that are secreted (count/pg insulin) in control and NOC-treated islets, immediately following a 12-hour radio-labeling process. Presented in (E) and (F) were (mean + SEM). The P values presented are from two-tailed type II t-test. “n”, the number of independent assays.

We next compared the secretion-probability of old and new IGs directly. Isolated islets were incubated overnight with 3H-labeled leucine/isoleucine and used in GSIS assays with or without NOC treatment. MT-disruption (in the presence of NOC) induced a significant increase in the proportion of older IGs being released (Figure 3F). These data support our model that the dense MT network in β cells traps older IGs. Without MTs, a bigger portion of older IGs will be made available for GSIS, leading us to explore the mechanisms that regulate both the MT networks and older vesicle docking/secretion in β cells. Note that the half-life of IGs in β cells was reported to be 3 – 5 days, we therefore consider IGs synthesized within the 12-hour window immediately before secretion assay as new IGs (25). This time-period allows sufficient amount of 3H-leucine/isoleucine incorporation in insulin for quantification.

### Gαo inactivation preferentially increases the secretion-probability of old IGs

The increased vesicle docking in MT-destabilized β cells is similar to what we have observed in the pancreatic specific *Gαo* mutant (*Gαo^F/F^; Pdx1^Cre^*) mouse β cells (15). Based on the published findings that trimeric G proteins can regulate MT dynamics (57, 65, 66), we explored the possibility that Gαo regulates IG secretion through MTs. We first tested if Gαo inactivation would impact the preferential secretion of newer IGs. As in the case of MT-destabilization, the *Gαo^F/F^; Pdx1^Cre^* β cells, wherein *Gαo* is efficiently inactivated in β cells (Figure 4A-D), secrete a larger portion of older vesicles under high-glucose stimulation (Figure 4E).

**Fig. 4.**
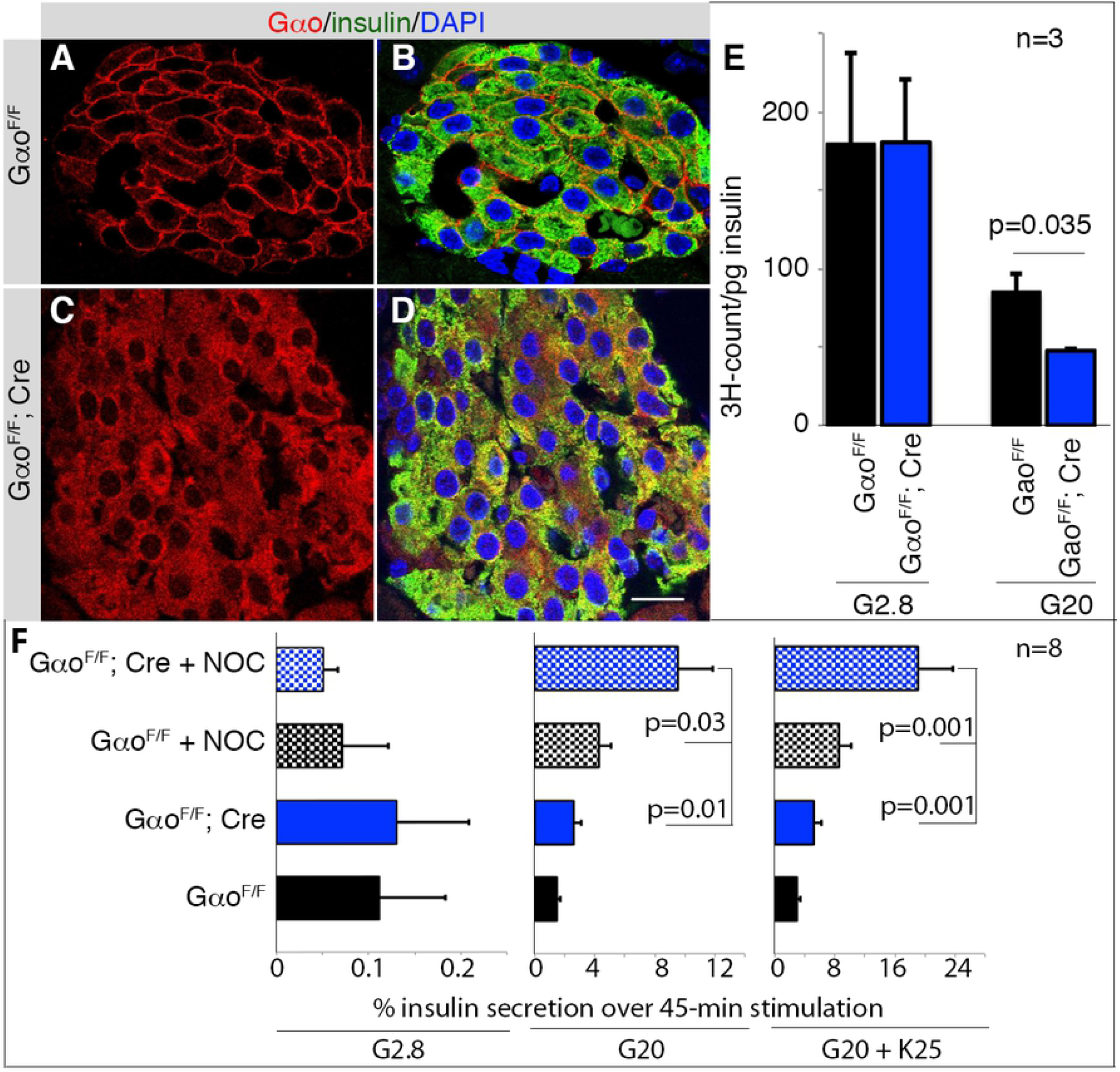
Gαo and MT regulate IG secretion via parallel pathways. (A-D) Immunoassays showing the complete Gαo inactivation in *Gαo^F/F^; Pdx1^Cre^* islet cells. Note that Cre-mediated *Gαo^F^* deletion will yield a mRNA that translates a short N-terminal Gαo peptide, recognized by the antibody (red) but has no detectable biological effect (15). Full length funtional Gαo is membrane-bound, while the N-terminal fragment is cytoplasmic, allowing for ready verification of *Pdx1^Cre^*-mediated *Gαo* inactivation in insulin+ (green) cells. DAPI (blue) stained for nuclei. Scale bar, 20 μm. (E) The levels of ^3^H-labeling in secreted insulin from control and *Gαo^F/F^; Pdx1^Cre^* β cells. (F) Insulin secretion results in *Gαo^F/F^; Pdx1^Cre^* islets, with or without NOC-treatment, induced by basal G2.8, stimulating G20, and [(A, B) G20 + K25 (25 mM KCl)]. Presented data in E and F are (mean + SEM). P values, results from two-tailed type II t-test.

### Gαo inactivation and MT disassembly independently regulate GSIS

We then tested if the increased GSIS in *Gαo^F/F^; Pdx1^Cre^* and MT-disrupted β cells depends on mobilizing a same pool of IGs. If so, we expect that MT disassembly in *Gαo^F/F^; Pdx1^Cre^* β cells will not further increase GSIS. In contrast, treating Gαo^-/-^ β cells with NOC induced an additional enhancement in GSIS (Figure 4F). Thus, different pools of IGs were mobilized/secreted in response to Gαo inactivation and MT disassembly, implying that Gαo does not directly regulate MTs in β cells.

### Gαo does not regulate MTs in β cells

We finally compared the MT density and stability in control and *Gαo^F/F^; Pdx1^Cre^* β cells via immunofluorescence. No differences in MT density (stained for tubulin) were observed between control and *Gαo^-/-^* β cells when examined in single cells (Figure 5A, C) or whole islets (Figure 5B, D), as measured by the average distances between MT filaments (Figure 5E). Similarly, we did not observe significant differences in MT-stability in *Gαo^F/F^; Pdx1^Cre^* and control β cells when the levels of Glu-tubulin [a well-established marker for MT stability, resulting from the removal of the C-terminal tyrosine in α tubulin to expose the glutamate residue (59)] were compared (Figure 5F-H). These findings suggest that inactivating Gαo does not alter MT dynamics in β cells, although both regulate IG docking and release.

**Fig. 5.**
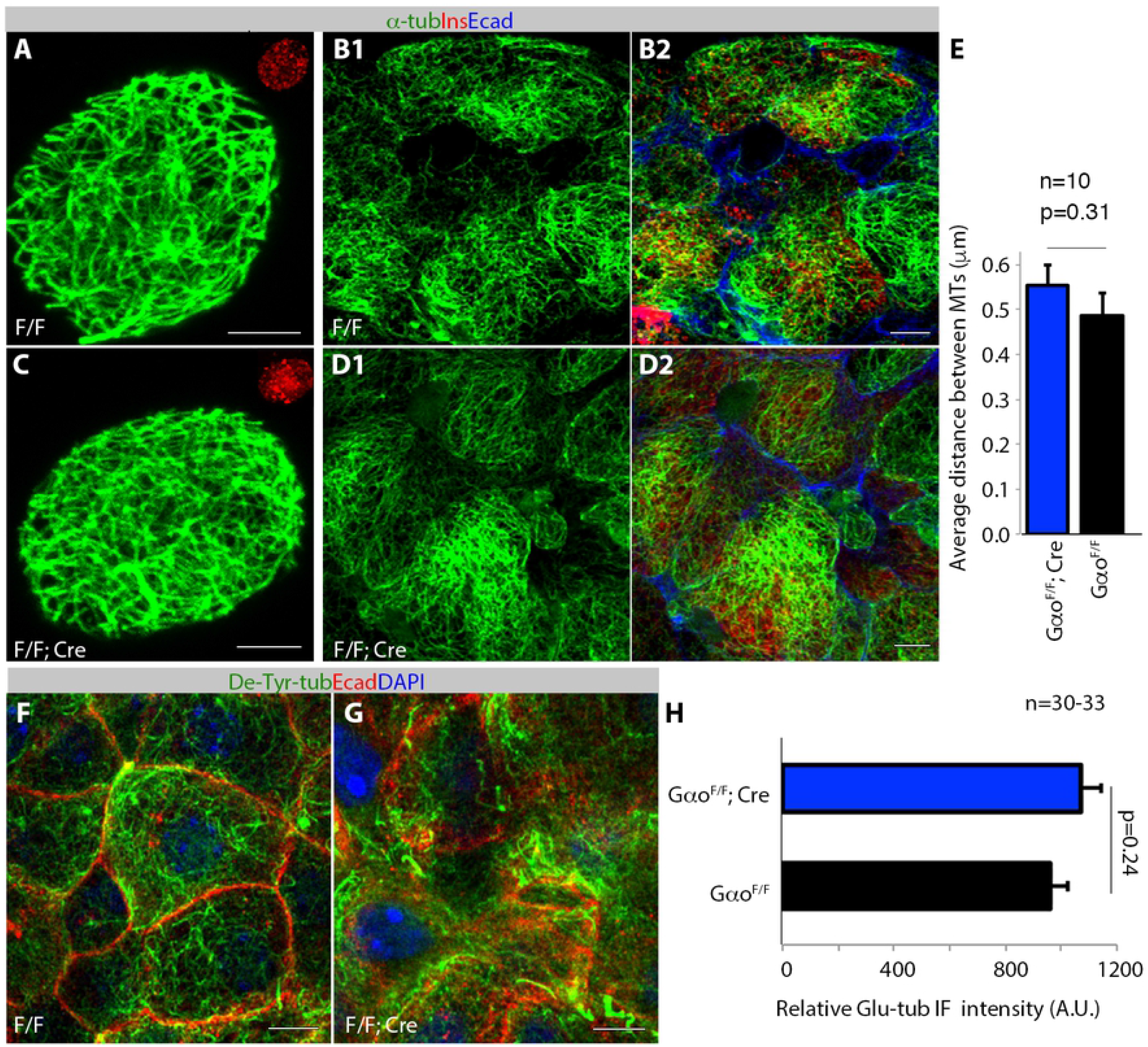
Inactivating *Gαo* does not alter MT dynamics in β cells. (A-E) The MT density in control and *Gαo^F/F^; Pdx1^Cre^* β cells, stained with anti-tubulin (green), anti-insulin (red), and anti-E-cadherin (blue) antibodies. Single cells attacehd to coverslips (A, C) or intact islets (B, D) were used. Insets in A, C, insulin staining to identify β-cells. Note that in (B, D), a single α-tubulin and a composite channel are presetented. The quantification data in E is MT density in single β cells, assayed as in (35). (F-H) The density of Glu-tubulin in control and *Gαo^F/F^; Pdx1^Cre^* β cells. Anti-Glu-tubulin (green), E-Cadherin (red), and DAPI (blue) were used. Presented data in E and H are (mean + SEM). P values, results from two-tail type II t-test. Scale bars, 5 μm.

In summary, we revealed two factors that act in parallel to regulate the preferential secretion of newly synthesized IGs in islet β cells. One is the MT, which originates from the Golgi and forms a non-directional meshwork. These properties makes these MTs ideal tracks for MT-dependent transport of newly produced IGs out of the trans-Golgi surface (40). However, they are unsuitable for directional bulk flow of IGs (35). As a result, the MTs act as holding places for IGs in the RP, whose transition to the RRP was expedited by both kinesin and dynein motor proteins (36, 38, 39). This mechanism is advantageous in that it prevents insulin over-secretion. However, when IGs age and lose their association with MTs, they will no longer be useful for function in the presence of a dense mesh of MTs and therefore must be degraded. This poises extra stress for β cells to replace this stock via insulin biosynthesis (67, 68). Note that although the absence of MTs can improve the usage of the old vesicles, β cells without MTs cannot maintain long-term function, due to the essential roles of MTs in β cells for new insulin biosynthesis (40).

The other factor that reduces the probability of secretion of old IGs is Gαo, whose inactivation enhances GSIS by increasing IG release of relatively older IGs as with MT-destabilization (15). Yet, Gαo does not regulate the MT stability or density in β cells as predicted based on *in vitro* biochemical studies (69, 70). An interesting future investigation could be to test if Gαo can regulate the affinity/processivity of motor proteins along the MTs. It would also be worthy to examine if Gαo regulates the affinity between IG and PM components that form the SNARE complex. The latter possibility is particularly attractive because Gαo has been shown to interact with Syntaxins (71), limiting factors for IG docking in β cells (52).

## Research Design and Methods

### Mice

Mouse usage followed protocols approved by the Vanderbilt University IACUC for GG/IK. Mice were euthanized by isoflurane inhalation. Wild type CD-1 (ICR) mice were from Charles River (Wilmington, MA). Production of Gαo^F/F^ and *Pdx1^Cre^* mice were described in (15), which were used to produce Gαo^-/-^ mutant β cells (*Gαo^F/F^; Pdx1^Cre^*).

### Islet isolation and routine GSIS

Islets were isolated from 8-16 week-old mice using collagenase perfusion as in (26). Briefly, ~2ml of 0.5 mg/mL of collagenase IV (Sigma, St. Louis, MI) dissolved in Hank’s Balanced Salt Solution (Corning, Corning, NY) was injected into the pancreas through the main duct. The pancreas was digested at 37°C for 20 minutes and washed 4 times with [RPMI-1640 media with 5.6 mM glucose (Gibco, Dublin, Ireland) + 10% heat inactivated (HI) fetal bovine serum (FBS, Atlanta Biologicals, Flowery Branch, GA)]. Islets were handpicked in the same media and let recover at 37 °C for at least 2 hours before down-stream experimentation.

GSIS follows routine procedures. Briefly, islets were washed twice with basal KRB solution (111 mM NaCl, 4.8 mM KC, 1.2 mM MgSO_4_, 1.2 mM KH_2_PO^4^, 25 mM NaHCO_3_, 10 mM HEPES, 2.8 mM glucose, 2.3 mM CaCl_2_, and 0.2% BSA). Islets were then incubated in the same solution (37 °C) for one hour, changed to new KRB to start the secretion assay. For insulin secretion induction, glucose (0.5 M) and/or KCl (1M) stock solutions were directly added to the KRB to desired concentrations. The secretion period assayed lasts 45 minutes. After secretion, islets were immediately frozen-thawed twice between −80 °C and room temperature. Acid alcohol extraction (70% alcohol + 0.2% HCl) was then performed at 4 °C overnight to determine the total insulin content. For each GSIS assays, 8-15 islets were used in 1 ml KRB. The insulin levels were then assayed using an ultrasensitive Insulin Elisa kit from Alpco after dilution to within the range of sensitivity.

### Islet pretreatment – MT destabilization, protein synthesis inhibition, and radiolabeling

For NOC treatment, islets were pre-incubated in KRB with 10 μg/ml NOC (with 20 mg/ml stock in DMSO) for one-hour to depolymerize MTs. This level of NOC was included in all solutions afterwards. DMSO (<0.05%) treated islets were used as controls.

To inhibit protein translation, islets were incubated in RPMI-640 with 10 mM glucose and 10 μM CHX for three hours to minimize the reduction of other proteins that are essential for secretion (63). Secretion assays follow as above in amino acid-free KRB for followup assays.

For radio-labeling, isolated islets were incubated for 12 hours in RPMI-1640 [supplemented with 10% FBS, 10 mM glucose, and 1/30 volume of ^3^H-labeled leucine/isoleucine (#NET1166001MC, Perkin Elmer)] cultured at 37°C with 5% CO_2_. Islets were then washed three times and chased in the same media without radioactive amino acids for 2 hours. GSIS assays were then performed as described above. In this case, the insulin levels were quantified using an Elisa kit. The radioactivity in secreted insulin was quantified using a scintillation counter (Beckman LS System 6000TA) following immunoprecipitation using guinea pig anti-insulin and Protein-A beads (ThermoFisher).

### Transmission electron microscopy (TEM) detection of IG-docking

Islets were incubated in RPMI-1640 with 10% FBS plus 5.6 or 20 mM glucose with or without 10 μg/ml NOC for 12 hours. Islets were then fixed, sectioned, and imaged following routine TEM protocols as detailed in (13). To count the number of docked IGs, Image J was used to measure the length of β-cell membrane. The docked vesicles, with near direct contact with the PMs (<10 nm away) were counted. The IG density was counted in a similar fashion, except that β-cell cytoplasmic areas were selected and measured. Double-blind tests were used without identifying the treatment conditions first.

### Immunofluorescence (IF) and microscopy

For Gαo staining, routine frozen pancreatic sections were used (26). Briefly, adult pancreata from mice of desired genotype were dissected and fixed at 4 °C overnight in 4% paraformaldehyde. Tissues were washed in PBS three-times and prepared as frozen sections, followed by immunofluorescence staining using mouse anti-Gαo described in (15). Insulin co-staining was used to identify β-cells. DAPI co-staining was used to locate nuclei.

For insulin, E-cadherin, and tubulin staining, single cells or islets were used as shown in (13). For single cells, islets were partially dissociated with trypsin, washed, overlaid onto human fibronectin-coated coverslips, and cultured overnight in RMPI-1640 media with FBS and 5.6 or 20 mM glucose. IF staining was then performed according to the following: cells or islets were extracted with methanol at −20°C for 5 minutes to remove free tubulin, followed by fixation with 4% paraformaldehyde (PFA) for 1 (for single cells) - 4 (for islets) hours, routine permeabilization and staining (13). The primary antibodies used were: rabbit anti-α-Tubulin (Abcam, Cambridge, UK, #ab18251), purified mouse anti-E-Cadherin (BD, San Jose, CA, #610181), rabbit anti-Glu-tubulin (MilliporeSigma #AB3201), and guinea pig anti-insulin (Dako, Santa Clara, CA, #A0564). The mouse anti-Gαo antibody was described in (15). Secondary antibodies are from Jackson ImmunoResearch (Alexa Fluor^®^ 647 AffiniPure Donkey Anti-Guinea Pig IgG (H+L) (706-605-148); Alexa Fluor^®^ 488 AffiniPure Donkey Anti-Rabbit IgG (H+L) (711545-152), and Alexa Fluor^®^ 594 AffiniPure Donkey Anti-Mouse (705-585-003). The dilution of all antibodies is 1:1000. Z-stacked images were captured using Nikon Eclipse A1R laser scanning confocal microscope. For super-resolution, images were captured at 0.125 μm intervals with DeltaVision OMX SIM Imaging System (GE technology) using a 60x NA1.4 lens and processed according to the manufacturer’s instruction.

### IG and MT density quantification

To quantify the IG localization in β cells (Figure 1), representative confocal images were taken. A line-scan was then performed along the long-axis in the image to quantify the IF intensity from one point in a β-cell border to another point at the opposite border. The length of the line was set arbitrarily at 10 and the intensity of the IF was measured using Image J. For this purpose, the lines were drawn to avoid the nuclei, which are devoid of insulin IF.

MT density quantification using Image J follows the procedure described in (35). Briefly, super-resolution MT images were captured as above at multiple z-depth. Line scans were then used to detect the size of the spaces between MTs by selecting 10 lines across the cell center with a 30-degree interval. A custom image J macro was used to create line selections and obtain the intensity profile. The lengths of regions without signal within the intensity profile were considered spaces between MTs, with the means and SEM presented.

### Statistics

For all studies, at least two biological repeats and two technical repeats were included. Student t-test, two-tailed type II analysis, was used for comparisons between two groups of data. Two-way ANOVA with Holm-Sidak’s multiple comparisons were used when more than two groups of data were compared. A *p* value below 0.05 was considered significant.

## Resource Availability

The reagents generated during the current studies are available from the corresponding authors upon reasonable request.

## Author Contributions

I.K. and G.G. conceptualized the work and designed the study. G.G. also performed radiolabeling and immunoprecipitation. R.H. performed GSIS assays. R. H., X.Z., and K.H. to perform immunoassays, imaging, and quantification. M. Y. did vesicle quantification. All authors contributed to writing the manuscript.

## Acknowledgement

This work was supported by National Institutes of Health (NIH) (https://www.niddk.nih.gov/ and https://www.nigms.nih.gov/) grant R01-DK106228 (to I.K. and G.G.), RO1-DK125696 and RO1-DK065949 (to G.G.), R35-GM127098 and R01-GM078373 (to I.K.). K.H. was supported by a postdoctoral fellowship from Eli Lilly and Company (LIFA fellowship 0101420) (https://www.amcp.org/resource-center/group-resources/residents-fellows/fellowships/Eli-Lilly).

